# Dysfunction of an energy sensor NFE2L1 triggers uncontrollable AMPK signal and glucose metabolism reprogramming

**DOI:** 10.1101/2021.09.07.459348

**Authors:** Qiufang Yang, Wenshan Zhao, Yadi Xing, Peng Li, Xiaowen Zhou, Haoming Ning, Ranran Shi, Shanshan Gou, Yalan Chen, Wenjie Zhai, Yahong Wu, Guodong Li, Zhenzhen Chen, Yonggang Ren, Yanfeng Gao, Yiguo Zhang, Yuanming Qi, Lu Qiu

**Author notes:** Qiufang Yang, Wenshan Zhao and Yadi Xing have contributed equally to this work. Correspondence to Dr. Lu Qiu or Prof. Yuanming Qi. School of Life Sciences, Zhengzhou University, Zhengzhou 450001, China.

## Abstract

NFE2L1 (also called Nrf1) acts a core regulator of redox signaling and metabolism homeostasis, and thus its dysfunction results in multiple systemic metabolic diseases. However, the molecular mechanism(s) by which NFE2L1 regulates glycose and lipid metabolism is still elusive. Here, we found that the loss of NFE2L1 in human HepG2 cells led to a lethal phenotype upon glucose deprivation. The uptake of glucose was also affected by NFE2L1 deficiency. Further experiments unveiled that although the glycosylation of NFE2L1 was monitored through the glycolysis pathway, it enabled to sense the energy state and directly interacted with AMPK. These indicate that NFE2L1 can serve as a dual sensor and regulator of glucose homeostasis. In-depth sights into transcriptome, metabolome and seahorse data further unraveled that glucose metabolism was reprogrammed by disruption of NFE2L1, so as to aggravate the Warburg effect in NFE2L1-silenced hepatoma cells, along with the mitochondrial damage observed under the electron microscope. Collectively, these demonstrate that disfunction of NFE2L1 triggers the uncontrollable signaling by AMPK towards glucose metabolism reprogramming in the liver cancer development.

## 1. Introduction

Redox homeostasis is a necessary prerequisite for maintenance of physiological responses, and the imbalance of redox homeostasis leads to various chronic systemic diseases. Nuclear factor erythroid 2 like 1 (NFE2L1/Nrf1), a core member of the cap’n’collar basic-region leucine zipper (CNC-bZIP) family, plays a critical role in regulating redox homeostasis in eukaryotic cells. Conventional knockout of *NFE2L1* could induce strong oxidative stress injury and result in fetal death. Also, conditional knockout of *NFE2L1* can increase cellular reactive oxygen species (ROS) level [1–4].

Studies have revealed that NFE2L1 plays an important role in metabolic pathway and influences the development of metabolic diseases. Human GWAS study shows that the single nucleotide polymorphism re3764400 which located in the 5’-flanking regions of *NFE2L1* gene is associated with obesity [5]. Overexpression of *NFE2L1* led to weight loss in transgenic mice, accompanied by insulin resistance symptoms such as increased pancreatic islets, increased insulin secretion, and increased blood glucose levels, eventually leading to diabetes mellitus [6]. In mouse pancreatic β cells, specific knockout of NFE2L1 caused hyperinsulinemia and glucose intolerance, the early symptoms of type II diabetes [7]. In adipocytes, specific knockout of *NFE2L1* gene resulted in almost complete disappearance of subcutaneous adipose tissue in mice, accompanied by insulin resistance, adipocyte hypertrophy and other obesity-related Inflammation [8]. And knockout of *NFE2L1* gene in the liver quickly caused non-alcoholic steatohepatitis (NASH) in mice [2,9]. All these results indicated that *NFE2L1* gene is essential for maintaining the homeostasis of lipid/carbohydrate metabolism.

Interestingly, research has shown that glycosylation of NFE2L1 protein mediates the cleavage and nuclear entry of NFE2L1 [10]. O-glycosylation modification in the Neh6L domain of NFE2L1 can reduce the transcription activity of NFE2L1 protein [11]. Additionally, in HepG2 cells, deletion of NFE2L1 can cause glucose deprivation, which finally induces cell death [12]. These studies suggest that NFE2L1 may be a key factor for regulating glucose metabolism homeostasis by sensing glucose levels. Consistently, our previous research also showed that loss of NFE2L1 expression can disrupt AMP-activated protein kinase (AMPK) signaling pathway [13], the core pathway involved in regulating energy metabolism [14,15]. In addition, metformin (MET), an AMPK signaling activator and the first-line drug for the treatment of diabetes, could inhibit NFE2L1 in an AMPK-independent manner, while the NFE2L1 could disruptive the activation of AMPK signal by MET [13], indicating the upstream regulation effects of NFE2L1 on AMPK signal.

Previous studies have suggested that NFE2L1 is essential for maintaining the homeostasis of glucose and lipid metabolism. However, the molecular mechanism is still obscure. Here, we investigated the effects of NFE2L1 deficiency on the glucose metabolism process in HepG2 cells and the results showed the dual functions of NFE2L1 as the sensor of glucose level and the regulator of glucose metabolism. At the same time, we also identified the mechanism of NFE2L1 in regulating glucose metabolism homeostasis through the verification of the effects of NFE2L1 in regulating AMPK signal.

## 2. Materials and Methods

### 2.1. Cell lines, culture and transfection

The information of the cell lines used in this study was shown in Table S1. Cells were growing in DMEM supplemented with 10% (v/v) FBS. The experimental cells were transfected by using Lipofectamine® 3000 Transfection Kit for 8 h, and then allowed for recovery from transfection in the fresh medium. The information of kits were shown in Table S1.

### 2.2. Expression constructs and other plasmids

The expression constructs for NFE2L1 and LKB1 were made by cloning the target sequences from full-length CDS sequences of *NFE2L1* and *LKB1* into *pLVX-EGFP-puro* vector. The primers used for these expression constructs were shown in Table S1.

### 2.3. Establishment of lentiviral infected cell lines

Lentiviruses for infection of HepG2^EGFP^ and HepG2^LKB1^ cells were packaged in HEK293T cells. HEK293T cells (5×10^5^) were seeded in 6-well plates and were transfected with three plasmids (1 μg of *pMD2.G*, 2 μg of *psPAX2*, 3 μg of either *pEGFP* or *pLKB1::EGFP* in 1mL of transfection volume) when cell confluence reached to 80%. The medium was collected and filtered by using 0.45 μm sterile filter to obtain the virus after 36~48 h. The virus was employed to transduce target cells, and puromycin (100 μM) was used to screen out the infected cells.

### 2.4. Cell viability assay

Experimental cells seeded in 96-well plates were processed according to the experimental design. The cytotoxic effects of indicated compound on experimental cells were determined by Cell Counting Kit-8 (CCK8) [16]. And the absorbance at 450 nm was measured by a microplate reader (SpectraMax iD5, USA). The information of kits were shown in Table S1.

### 2.5. Cellular ROS staining

Experimental cells were allowed for growing to reach an appropriate confluence in 6-well plates and then incubated in a serum-free medium containing 10 μM of 2′,7′-Dichlorodihydrofluorescein diacetate (DCFH-DA) [17] at 37□ for 20 min. Thereafter, the cells were washed three times with serum-free media, before the green fluorescent images were achieved by microscopy.

### 2.6. Glucose uptake assay

The uptake of 2-Deoxy-2-[(7-nitro-2,1,3-benzoxadiazol-4-yl)amino]-D-glucose (2-NBDG), which is a fluorescent glucose analog, is used to visualizing the uptake capacity of glucose by living cells. Experimental cells were allowed for incubated in a serum-free medium containing 20 μM of 2-NBDG at 37□ for 10 min. Thereafter, the cells were washed three times with serum-free media, before the green fluorescent images were achieved by microscopy.

### 2.7. RNA isolation and real-time quantitative PCR (qPCR)

Experimental cells were subjected to isolate total RNAs using the RNAsimple Kit. Then, 1 μg total RNAs were added in a reverse-transcriptase reaction to generate the first strand of cDNA, by using the RevertAid First Strand Synthesis Kit. The synthesized cDNA served as the template for qPCR with distinct primers, which occurred in the GoTaq® qPCR Master Mix. The mRNA expression level of β-actin was selected as an optimal internal standard control. The primers used for qPCR and the information of kits were shown in Table S1.

### 2.8. Western bloting (WB), Co-Immunoprecipitation (Co-IP) and protein deglycosylation reactions

The total protein extraction buffer and WB were carried out as previously described [13]. The Co-IP were carried out as previously described [18]. Before visualized by WB, the deglycosylation reactions of samples with Endo H in vitro were carried out as described previously [19]. The information of all antibodies used herein were shown in Table S1.

### 2.9. Transcriptome sequencing

The cultured cells (HepG2^shNC^ and HepG2^shNFE2L1^) were lysed by Trizol, and the transcriptome sequencing by *Beijing Genomics Institute (BGI)* with the Order Number is F21FTSCCKF2599_HOMdynrN. The samples were measured use the BGISEQ-500 platform. The transcriptome sequencing results are shown in Table S2.

### 2.10. Metabolomics testing

The HepG2^shNC^ and HepG2^shNFE2L1^ cells were digested and counted, and the cell pellets were obtained by low-speed centrifugation. The cell samples were immediately frozen in liquid nitrogen. Metabolomics testing by *Metabolomics and Systems Biology Company, Germany*. And the Order Number is POCMTS2016006. The samples were measured with a Waters ACQUITY Reversed Phase Ultra Performance Liquid Chromatography (RP-UPLC) coupled to a Thermo-Fisher Exactive mass spectrometer. For each type of cells, three replicates were extracted and measured.

### 2.11. Electron Microscopy

The adherent cells were digested with trypsin and fixed with 2.5% glutaraldehyde. The ultrastructure of cells was examined by transmission electron microscope (JEM-HT7700 model, Hitachi, Japan) with 5um, 1um, and 500nm scales, respectively.

### 2.12. Seahorse Metabolic Analyzer

The oxygen consumption rate (OCR) of cells was measured by Seahorse XF Cell Mito Stress Test Kit. The experimental operation was carried out according to the instructions. The OCR were measured using the Seahorse Metabolic Analyzer to characterize the cell’s oxidative phosphorylation level. The information of Kits were shown in Table S1.

### 2.13. Assays of ATP and Lactate Levels in cells

The ATP levels and Lactate levels of cells are determined according to the instruction of ATP assay kit and Lactic Acid assay kit. The experimental operation was carried out according to the instructions. The information of Kits were shown in Table S1.

### 2.14. Statistical analysis

The data were here presented as a fold change (means ± S.D.), each of which represents at least three independent experiments that performed in triplicate. Significant differences were statistically determined by either the Student’s t-test or Multiple Analysis of Variations (MANOVA). The statistical significance was defined by symbols ‘*’ means *p* < 0.05, whilst the ‘*n.s.*’ letters represent ‘no significant’.

## 3. Results

### 3.1. NFE2L1 deficiency disrupts cellular energy metabolism signals and induces cell death by glucose starvation

Previous studies have shown that mice with NFE2L1 overexpression or knockout exhibit diabetes-like phenotypes [6–8]. Metformin has shown regulatory effects on NFE2L1 [13]. These studies indicated the involvement of NFE2L1 in maintaining the homeostasis of glucose metabolism. To further explore the relationship between NFE2L1 and glucose metabolism, HepG2 cell line with *NFE2L1* knockdown was constructed (Figure 1A). Through the glucose deprivation experiment, HepG2 cells in the medium without glucose lacking *NFE2L1* expression were more susceptible to death (Figure 1B,C). Glucose is the major source of energy for cells, but it is not an essential nutrient. Here, WZB117 [20], a glucose uptake inhibitor, was used to test the effects of impaired glucose uptake on cells. The results showed that compared with the control group, the *NFE2L1* knockdown cells was exhibited enhanced killing activity by treating with WZB117 (Figure 1D), which indicate that loss of NFE2L1 may disrupt the homeostasis of energy metabolism.

**Figure 1.**
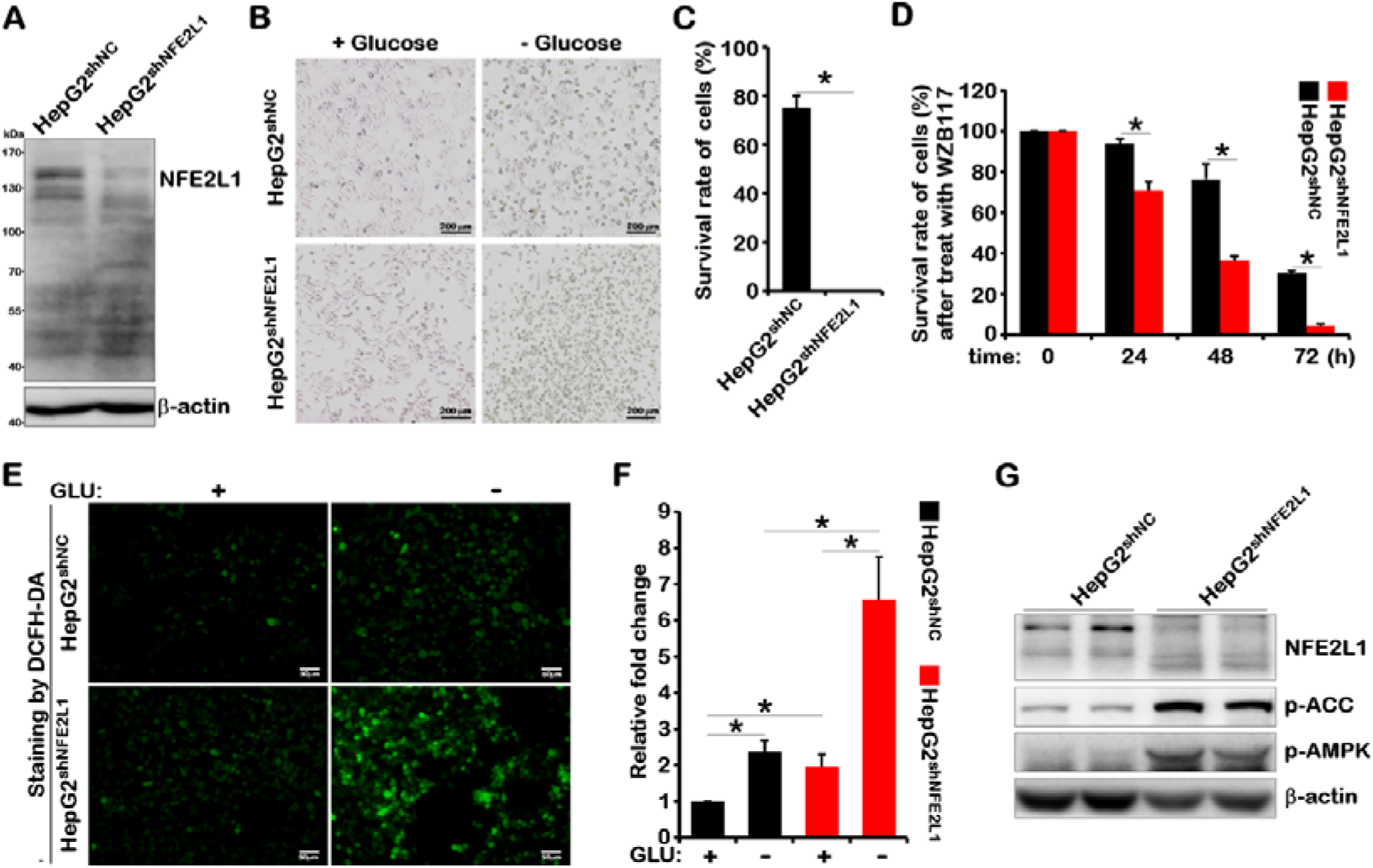
NFE2L1 knockdown caused HepG2 cells to be more sensitive to glucose starvation. (**A)** The total protein of HepG2^shNC^ and HepG2^shNFE2L1^ cells were collected, then the expression of NFE2L1 and β-actin were detected by WB. The HepG2^shNC^ and HepG2^shNFE2L1^ cells were cultured used the DMEM with or without glucose (2 g/L or 0 g/L) for 18h, the morphology of cells were observed under the microscope **(B)**, and the survival rate of cells was detected by CCK8 kit **(C)**. (**F)** The HepG2^shNC^ and HepG2^shNFE2L1^ cells were treated with WZB117 (200μM), and the survival rate of cells was detected in 0h, 24h, 48h, 72h by CCK8 kit, respectively. The HepG2^shNC^ and HepG2^shNFE2L1^ cells were cultured used the DMEM without glucose for 6h, then the ROS in cells were staining by DCFH-DA (10μM) for 20min and achieved by microscopy (**E**), and the intensity of fluorescence were counted **(D)**, the scale is 50μm. (**G)** The total protein of HepG2^shNC^ and HepG2^shNFE2L1^ cells were collected, then the expression of NFE2L1, pAMPK, pACC and β-actin were detected by WB. n ≥ 3, ‘*’ means *p* < 0.05.

NFE2L1 is a regulator of redox homeostasis, and most of the phenotypes related to NFE2L1 are related to oxidative stress. Results showed that the ROS level in the control group (HepG2^shNC^) was increased by about 2 times when the glucose was deprived for 6 hours. In *NFE2L1* knockdown cells (HepG2^shNFE2L1^), the level of ROS was increased by about 3 times, and the shrinking of cells has been observed (Figure 1E,F). In *NFE2L1* knockdown cells, the ROS level was increased more than 6 times in comparison with the control group after glucose deprivation for 6 hours, indicating the high levels of oxidative damage of *NFE2L1* knockdown cells. These results were in accordance with previous studies that glucose starvation resulted in strong oxidative stress to induce cell death after *NFE2L1* knockout [3,12].

AMPK, the core signal of energy metabolism, is the guardian of metabolism and mitochondrial homeostasis [21]. Western bloting (WB) results showed knockdown of *NFE2L1* caused a significant increase in the phosphorylation level of AMPK and its downstream protein acetyl-CoA carboxylase alpha (ACC) [22] (Figure 1G), indicating that the cells may be in a starvation state even in the sufficient nutritional conditions. This phenomenon may due to the lack of NFE2L1 in HepG2^shNFE2L1^ cells. All these results suggested that NFE2L1 could function through sensing the energy state of the cells.

### 3.2. Glycolysis promotes glycation of NFE2L1 and lack of NFE2L1 promotes glucose absorption

In the cell culture system, in addition to glucose, other nutrients also contribute to provide the beneficial environment for cell growth, including cytokines and fetal bovine serum (FBS). Studies have shown that NFE2L1 could be activated by FBS/mechanistic target of rapamycin kinase (mTOR)/sterol regulatory element binding transcription factor 1 (SREBF1) pathway [23]. The mTOR and AMPK signaling pathways could be affected by NFE2L1 overexpression or knockdown [13], indicating the correlation of NFE2L1 with energy metabolism. Therefore, the alternation of NFE2L1 in response to the change of glucose and serum concentration was tested in the cell culture medium and the results showed that addition of FBS could upregulate the total protein content of NFE2L1, and glucose could directly affect the bands of NFE2L1 (Figure 2A,B). The level of p-AMPK gradually decreased with the increase of serum concentration in the presence and absence of glucose (Figure 2A). And the increased glucose concentration slightly altered the p-AMPK level with and without FBS (Figure 2B). The p-mTOR level could be significantly elevated with the presence of glucose (Figure 2A) and FBS (Figure 2B) despite the concentration alternation of glucose and FBS.

**Figure 2.**
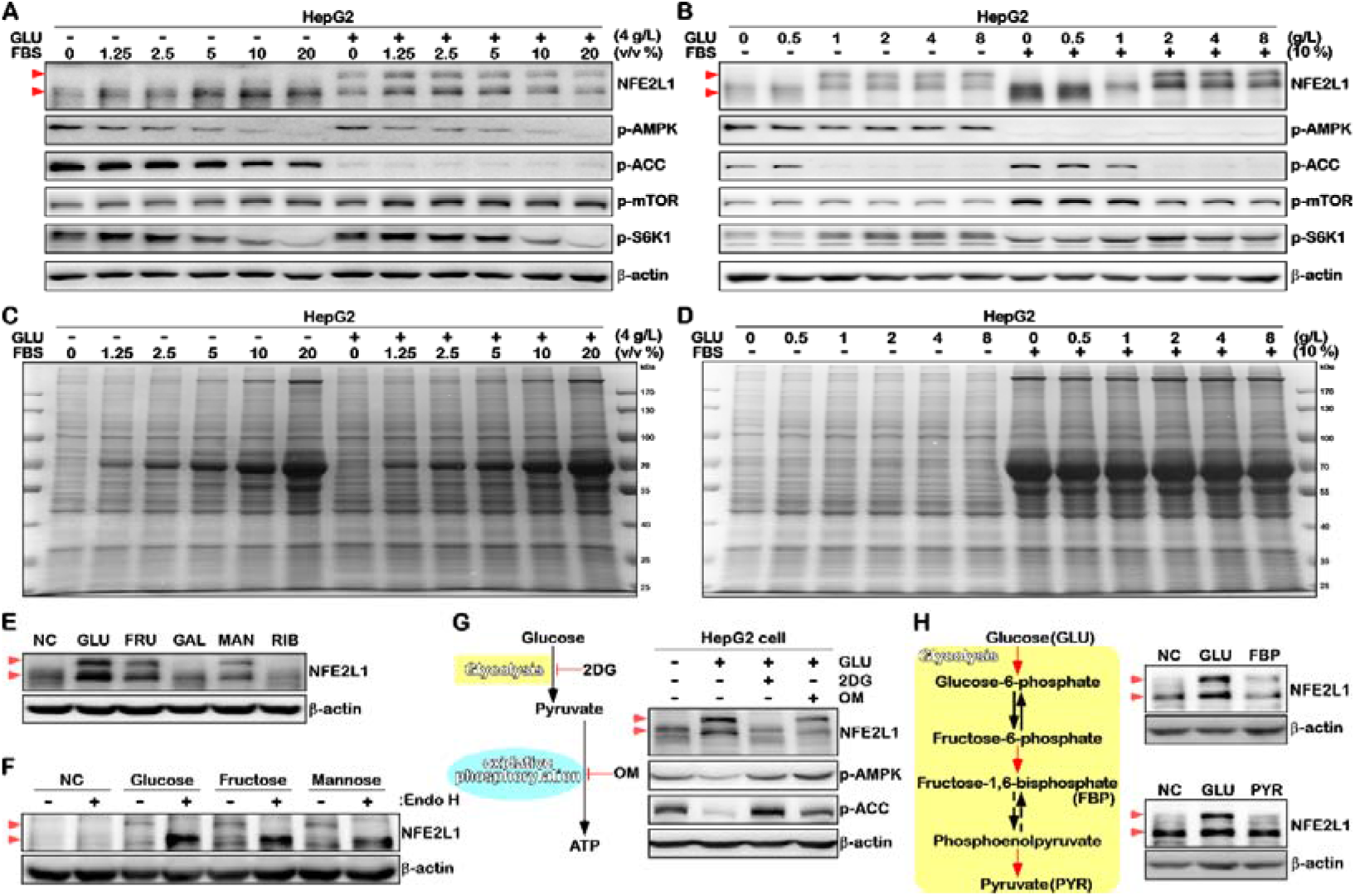
NFE2L1 is a glucose sensitive protein. **(A)** HepG2 cells were cultured used DMEM medium with different volume ratios of FBS (0%, 1.25%, 2.5%, 5%, 10%, 20%), with or without glucose (4g/L or 0g/L) for 16h, and the total protein were collected, then the expression of NFE2L1, p-AMPK, p-ACC, p-mTOR, p-S6K1 and β-actin were detected by WB. (**B)** HepG2 cells were cultured used DMEM medium with different concentrations of glucose (0g/L, 0.5g/L, 1g/L, 2g/L, 4g/L, 8g/L), and with or without FBS (10% or 0%) for 16h, and the total protein were collected, then the expression of NFE2L1, p-AMPK, p-ACC, p-mTOR, p-S6K1 and β-actin were detected by WB. (**C)** The total protein change of the sample in (**A**) staining by coomassie brilliant blue. (**D)** The total protein change of the sample in (**B**) staining by coomassie brilliant blue. (**E)** HepG2 cells were glucose starvation for 4h and then cultured used DMEM with GLU (2g/L), FRU (30mM), GAL (30mM), MAN (30mM) and RIB (30mM) for 4h, the total protein were collected and the expression of NFE2L1 and β-actin were detected by WB. (**F)** The HepG2 cells were cultured used DMEM medium with GLU (0g/L), GLU (2g/L), FRU (30mM) or MAN (30mM) for 4h, the total protein were collected and Endo H were used to deglycosylate the glycated protein in vitro, then the expression of NFE2L1 and β-actin were detected by WB. (**G)** The HepG2 cells were glucose starvation for 4h and then treated with GLU (2g/L), GLU (2g/L) + 2DG (20mM) or GLU (2g/L) + OM (10μM), and the total protein was collected after 4 hours, then the expression of NFE2L1, p-AMPK, p-ACC, p-mTOR, p-S6K1 and β-actin were detected by WB. (**H)** The HepG2 cells were glucose starvation for 4h and then treated with GLU (2g/L), FBP (30mM), or PYR (30mM) for 4h, the total protein was collected and then the expression of NFE2L1 and β-actin were detected by WB.

Activation of AMPK and mTOR signals can be characterized by p-ACC and phosphorylated- ribosomal protein S6 kinase B1 (p-S6K1) expression respectively. In fact, as the results showed, p-AMPK was more sensitive to FBS in HepG2 cells (Figure 2B), while p-ACC was more sensitive to glucose (Figure 2A). The expression level of p-S6K1 increased at low concentrations of FBS and decreased when exposed to higher concentrations of FBS (Figure 2A,B). With the increased concentration of glucose, the level of p-S6K1 was upregulated (Figure 2A,B). In addition, the bands of p-AMPK and p-S6K1 moved down with the increase of FBS, which might be due to the fact that the protein synthesis upregulated in respond to the increased amount of FBS, as the results showed by coomassie brilliant blue staining of protein samples in Figure 2C and Figure 2D. It is important to note that the level of p-ACC has a strong negative relationship with the glycation of NFE2L1 (Figure 2A,B). Combined with the increase of p-ACC expression after NFE2L1 knockdown (Figure 1G) and the inhibition of p-ACC by NFE2L1 overexpression, these results together suggested that NFE2L1 might directly affect the catalytic activity of p-AMPK on substrates.

Studies have shown that glycosylation can regulate the activity of NFE2L1 [10,11,24,25]. Here, the effects of five common monosaccharides on NFE2L1 protein were tested and the results indicated that fructose (FRU) and mannose (MAN), similar as glucose (GLU), could induce the production of NFE2L1 with larger molecular weight (Figure 2E), which could be ascribed to the glycosylation of NFE2L1 protein as it revealed by the de-glycosylation experiment shown in Figure 2F. On the contrary, galactose (GAL) and ribose (RIB) showed null effects on the molecular weight of NFE2L1 protein (Figure 2F).

The catabolism of carbohydrates mainly undergoes anaerobic glycolysis in the cytoplasm and aerobic oxidation in the mitochondria. In HepG2 cells, treatment with 2-Deoxy-D-glucose (2DG, a glycolysis inhibitor) [26] could effectively inhibit glycolysis of NFE2L1. Inhibition of oxidative phosphorylation process using Oligomycin (OM) [27] could decrease the total expression level of NFE2L1 (Figure 2G). Further results showed that fructose-1,6-bisphosphate (FBP), the intermediate product of the glycolysis process, induced the glycation of NFE2L1, while the pyruvate (PYR) had no such effects (Figure 2H). These findings implied that activation of hexokinase (HK) and phosphofructokinase, liver (PFKL), the key rate-limiting enzymes of glycolysis, or inhibition of pyruvate kinase M1/2 (PKM) may have the function of activating NFE2L1.

The above results demonstrated that NFE2L1 was a glucose-sensitive protein. As the results revealed in Figure 1, knockdown of NFE2L1 induced cell death by glucose starvation and activation of AMPK signaling, indicating that NFE2L1 might be involved in feedback regulation of energy metabolism. Our previous studies have shown that NFE2L1 could be inhibited by metformin (MET), and knockdown of NFE2L1 disrupted the activation of AMPK signal by MET. In addition, studies have shown that MET promoted the uptake of glucose in HepG2 cells [28]. These studies suggested the promotion of MET on glucose uptake in HepG2 cells might be related to its inhibition of NFE2L1.

NFE2L1 deficiency improved the glucose uptake in HepG2 cells. Interestingly, knockdown of NFE2L1 counteracted the promotion of MET on glucose uptake of HepG2 cells (Figure 3A,B). Similarly, loss of NFE2L1 expression also neutralized the ROS elevation induced by MET [13]. These data indicated that the inhibitory effect of MET on NFE2L1 might be the main reason for the improvement of the glucose uptake of HepG2 cells. However, it could not be ruled out that the glucose uptake capacity of cells reached a threshold. Subsequently, the changes of glucose transporter genes in transcriptome date were checked, and the data showed that the *solute carrier family 2 member 1* (*SLC2A1/GLUT1*) and *SLC2A13* were increased, while the *SLC2A3*, *SLC2A6* and *SLC2A8* were depressed after *NFE2L1* knockdown (Figure 3C). And the qPCR results showed that *SLC2A3* was significantly reduced, while *SLC2A4* and *SLC2A6* were obviously increased (Figure 3D). The *SLC2A2*, *SLC2A5*, *SLC2A7* and *SLC2A9* genes were not detected by real-time quantitative PCR (qPCR), which might be due to the low or lack of expression of these genes. By comparing the transcriptome and qPCR results, it was found that only *SLC2A3* was consistently reduced, but it was opposite of the phenotype. Then, the protein levels of glucose transporter were been detected and the results showed that the protein expression level of GLUT3 was significantly increased after NFE2L1 knockdown (Figure 3E), indicating that the increase in glucose uptake caused by NFE2L1 knockdown might be related to GLUT3. In addition, the mRNA level of *SLC2A1*, of which the protein level could hardly been detected, was highest, suggesting that GLUT1 might be severely inhibited at the post-transcriptional level in HepG2 cells.

**Figure 3.**
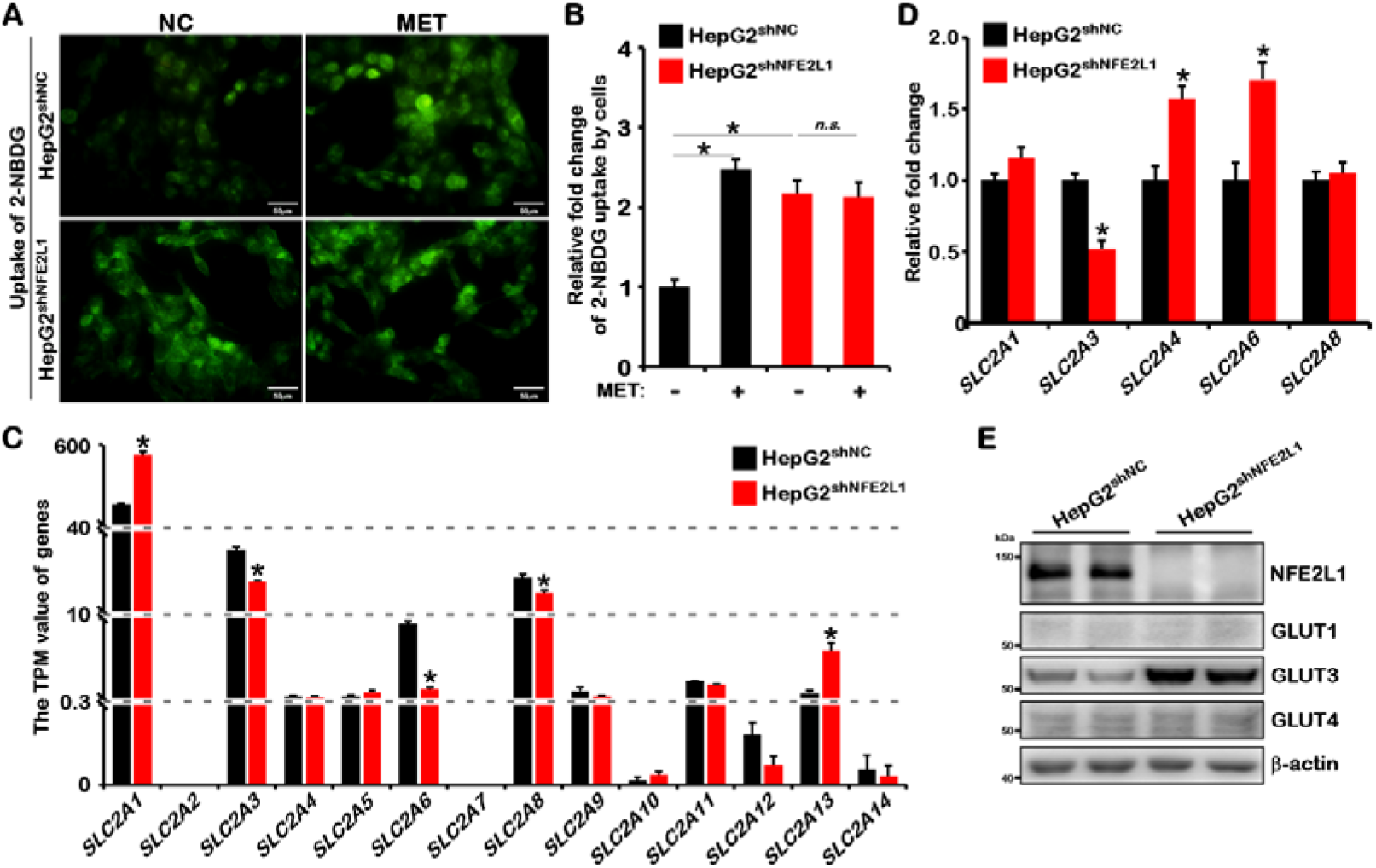
NFE2L1 can affect the uptake of glucose for HepG2 cells. **(A)** The HepG2^shNC^ and HepG2^shNFE2L1^ cells were treated with MET (1mM) for 12h, and then incubated in a serum-free medium containing 20μM of 2-NBDG at 37□ for 10min and the green fluorescent images were achieved by microscopy, the scale is 50μm. (**B)** The statistics show the results in (**A**). (**C)** The TPM value of *SLC2A1*, *SLC2A2*, *SLC2A3*, *SLC2A4*, *SLC2A5*, *SLC2A6*, *SLC2A7*, *SLC2A8*, *SLC2A9*, *SLC2A10*, *SLC2A11*, *SLC2A12*, *SLC2A13*, *SLC2A14* genes that related to glucose uptake of cell, date comes from transcriptome sequencing. (**D)** The expression of *SLC2A1-9* genes in HepG2^shNC^ and HepG2^shNFE2L1^ cells were detected by qPCR. The *SLC2A2*, *SLC2A5*, *SLC2A7* and *SLC2A9* have not be detected for it not expressed or extremely low, and β*-actin* gene was used as the internal control. (**E)** The expression of NFE2L1, GLUT1, GLUT3, GLUT4 and β-actin in HepG2^shNC^ and HepG2^shNFE2L1^ cells were detected by WB. n ≥ 3, ‘**\***’ means *p* < 0.05, ‘*n.s.*’ means ‘no significant’.

### 3.3. NFE2L1 deficiency results in the reprogramming of glucose metabolism and exacerbates the Warburg effect

The above results revealed the function of NFE2L1, as glucose metabolism sensor, to regulate glucose uptake. Next, the effects of NFE2L1 knockdown on the key rate-limiting enzymes implicated in glycolysis, gluconeogenesis and tricarboxylic acid (TCA) cycle were tested (Figure 4A). As the results shown in Figure 4B, increase of *heme oxygenase 1* (*HMOX1*) and *prostaglandin-endoperoxide synthase 2* (*PTGS2*/*COX2*), and decrease of *PTGS1* were detected accompanied by NFE2L1 knockdown, suggesting the loss of function of NFE2L1 [3]. *HK1*, the key enzyme involved in catalyzing the conversion of glucose to glucose-6-phosphate, was significantly increased. *PFKL* and phosphofructokinase, muscle (*PFKM*), which catalyze the conversion of fructose-6-phosphate to fructose-1,6-bisphosphate, was reduced. The expression of *PKM*, which has been involved in catalyzing the conversion of phosphoenolpyruvate to pyruvate, was increased. No apparent change of the key enzyme gene *dihydrolipoamide dehydrogenase* (*DLD*) that catalyzes the entry of pyruvate into the TCA cycle was detected. The expression of the *pyruvate carboxylase* (*PC*) gene was significantly reduced, indicating that the metabolic flow from glycolysis to the TCA cycle might be restricted (Figure 4C).

**Figure 4.**
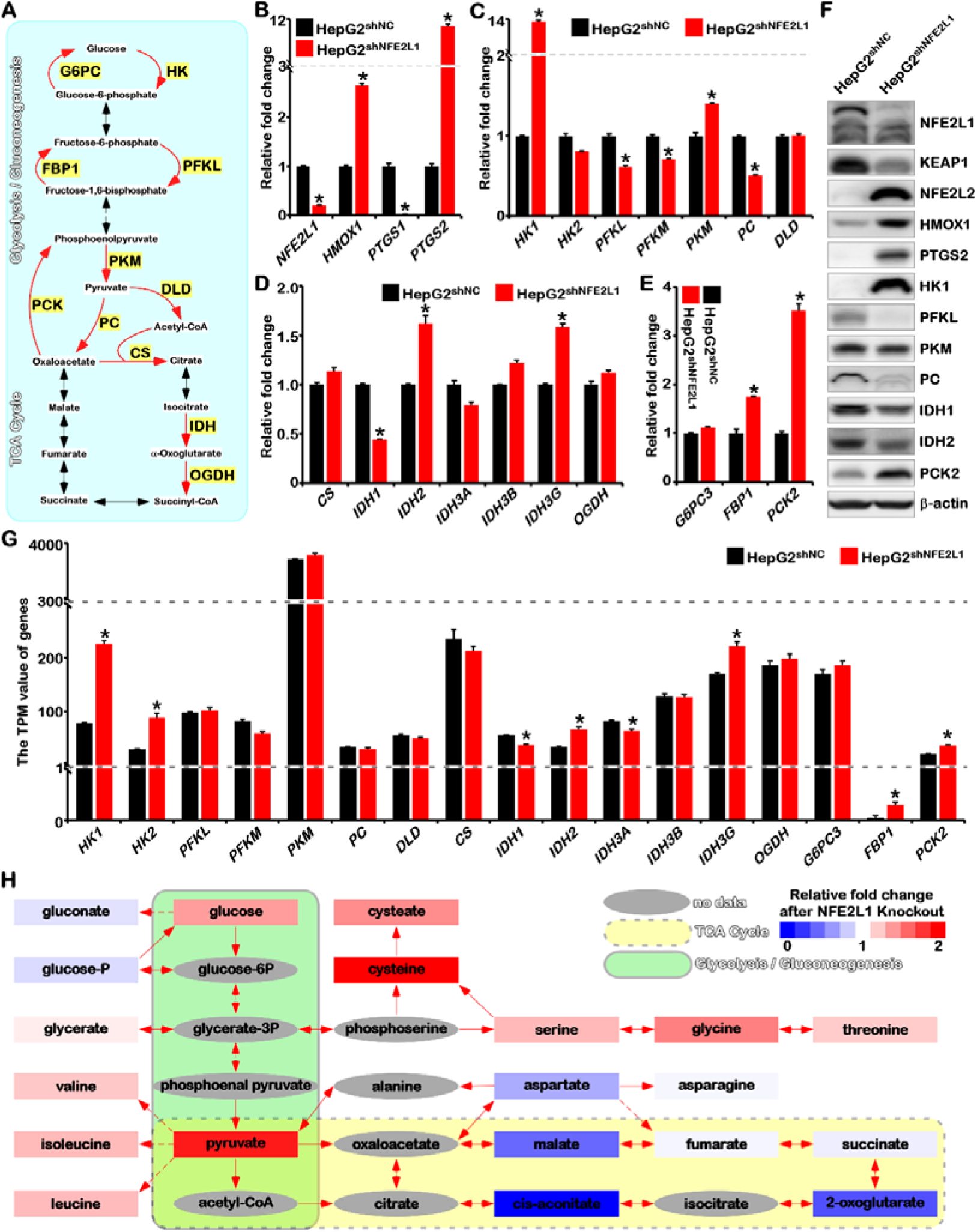
Knockdown of NFE2L1 leads to reprogramming of glucose metabolism. (**A)** The schematic diagram of glycolysis, gluconeogenesis and TCA metabolism process and key enzymes. (**B)** The expression of *NFE2L1*, *HMOX1*, *PTGS1* and *PTGS2* genes in HepG2^shNC^ and HepG2^shNFE2L1^ cells were detected by qPCR, β*-actin* gene was used as the internal control. **(C)** The expression of *HK1*, *HK2*, *PFKL*, *PFKM*, *PKM*, *PC* and *DLD* genes in HepG2^shNC^ and HepG2^shNFE2L1^ cells were detected by qPCR, β*-actin* gene was used as the internal control. **(D)** The expression of *CS*, *IDH1*, *IDH2*, *IDH3A*, *IDH3B*, *IDH3G* and *OGDH* genes in HepG2^shNC^ and HepG2^shNFE2L1^ cells were detected by qPCR, β*-actin* gene was used as the internal control. **(E)** The expression of *NFE2L1*, *G6PC3*, *FBP1* and *PCK2* genes in HepG2^shNC^ and HepG2^shNFE2L1^ cells were detected by qPCR, β*-actin* gene was used as the internal control. (**F)** The expression of NFE2L1, KEAP1, NFE2L2, HMOX1, PTGS2, HK1, PFKL, PKM, PC, IDH1, IDH2, PCK2 and β-actin in HepG2^shNC^ and HepG2^shNFE2L1^ cells were detected by WB. (**G)** The expression of key enzyme for glucose metabolism in the transcriptome data. (**H)** The results of the metabolome showed the effect of NFE2L1 on Central Metabolism. n ≥ 3, ‘*’ means *p* < 0.05.

Then the expression of key enzyme genes involved in TCA cycle was detected and the results showed that *citrate synthase* (*CS*), which catalyzes to produce citrate, and *oxoglutarate dehydrogenase* (*OGDH*), which catalyzes to produce succinyl-CoA, exhibited no significant changes in mRNA levels. And *isocitrate dehydrogenase (NADP(+)) 1* (*IDH1*), which catalyzes the conversion of isocitrate to a-oxoglutarare, was reduced. The level of *IDH2* and *IDH3G* was upregulated (Figure 4D). It can be inferred from these results that the TCA cycle displayed no remarkable changes. *phosphoenolpyruvate carboxykinase 2* (*PCK2*) and *fructose-bisphosphatase 1* (*FBP1*), the key rate-limiting enzymes in the gluconeogenesis process, were significantly increased, while no difference of *glucose-6-phosphatase catalytic subunit 3* (*G6PC3*) was detected (Figure 4E), indicating that the gluconeogenesis process might be enhanced as NFE2L1 absence.

The western blotting result showed that NFE2L1 knockdown could efficiently elevate the expression of NFE2L2, HMOX1, and PTGS2 and decrease kelch like ECH associated protein 1 (KEAP1) (Figure 4F), which were consistent with our previous results [3]. Then the changes in the protein levels of the key enzymes of sugar metabolism were detected. The WB results of these key proteins were almost the same as the gene expression (Figure 4F). The protein expression of IDH1 and IDH2 both decreased. Combined with the significant decrease of PC and the increase of PCK2, these results indicated the enhancement of the glycolysis and gluconeogenesis, while the process from anaerobic glycolysis to aerobic oxidation was been blockaded. Consistently, the transcriptome data also suggested that glycolysis and gluconeogenesis were enhanced after knockdown of NFE2L1 in HepG2 cells (Figure 4G).

In the process of glucose metabolism, loss of NFE2L1 may induce increased glycolysis and gluconeogenesis and the suppressive oxidative phosphorylation. These results indicated that NFE2L1 deficiency might trigger the Warburg effect, consistent with that specific knockout of *NFE2L1* in liver tissue could trigger the Non-alcoholic fatty liver disease (NAFLD) and hepatocellular carcinoma (HCC) in the mouse model [2]. Analysis of metabolomic data showed that, in the *NFE2L1* knockout HepG2 cells, the content of glucose metabolism intermediate products glucose and pyruvate were increased, and amino acids closely related to glycolysis intermediate products were significantly increased, such as glycerate, valine, isoleucine, leucine, cysteate, cysteine, serine, glycine, and threonine (Figure 4H). Malate, 2-oxoglutarate, and cis-aconitate, the TCA cycle intermediate products, were significantly reduced (Figure 4H). These results implied that the absence of NFE2L1 produced a phenotype similar to the warburg effect. The enhancement of warburg effect, one of the metabolic characteristics of tumor cells, often means the malignant transformation of tumors. It is worth mentioning that the metabolome data showed that the loss of NFE2L1 expression caused an increase in the levels of various amino acids in cells, which might be the reason for the activation of mTOR signal caused by NFE2L1 knockdown [13,29,30].

### 3.4. NFE2L1 knockdown causes the damage of mitochondrial function

Consistent with our results, previous studies had shown that overexpression and knockout of NFE2L1 could change glucose metabolism through different ways such as ROS, insulin secretion, and liver metabolism [6,7,12]. However, the underlying molecular mechanism by which NFE2L1 affects glucose metabolism remains to be explored. As a transcription factor, NFE2L1 could influence thousands of downstream genes with various functions [3,31,32], making it difficult to identify the detailed mechanism of NFE2L1 in affecting glucose metabolism. Actually, the effect of NFE2L1 deficiency on glucose metabolic reprogramming is likely to be a systemic response.

Previous result showed that the TCA cycle in the mitochondria was inhibited (Figure 4), which made it necessary to explore the effects of NFE2L1 on mitochondrial function. Under electron microscope, it was found that NFE2L1 knockdown significantly reduced the number of mitochondria. The remaining mitochondria caused by NFE2L1 knockdown possessed much smaller size and the ridge-like structure was significantly reduced (Figure 5A,B). In NFE2L1 knockdown cells, accumulation of lipid droplets could be found, which is consistent with the lipid accumulation induced by special knockout of NFE2L1 in mouse live [2]. In addition, more autophagosomes could be found in NFE2L1 knockdown cells (Figure 5A,B), suggesting that damaged mitochondria could be transferred to autophagic mitochondria to be degraded. Consistently, studies have shown that AMPK can promote mitochondrial division and mitochondrial autophagy [33,34], suggesting that the alternation of mitochondrial caused by NFE2L1 knockdown may be related to increased AMPK signals.

**Figure 5.**
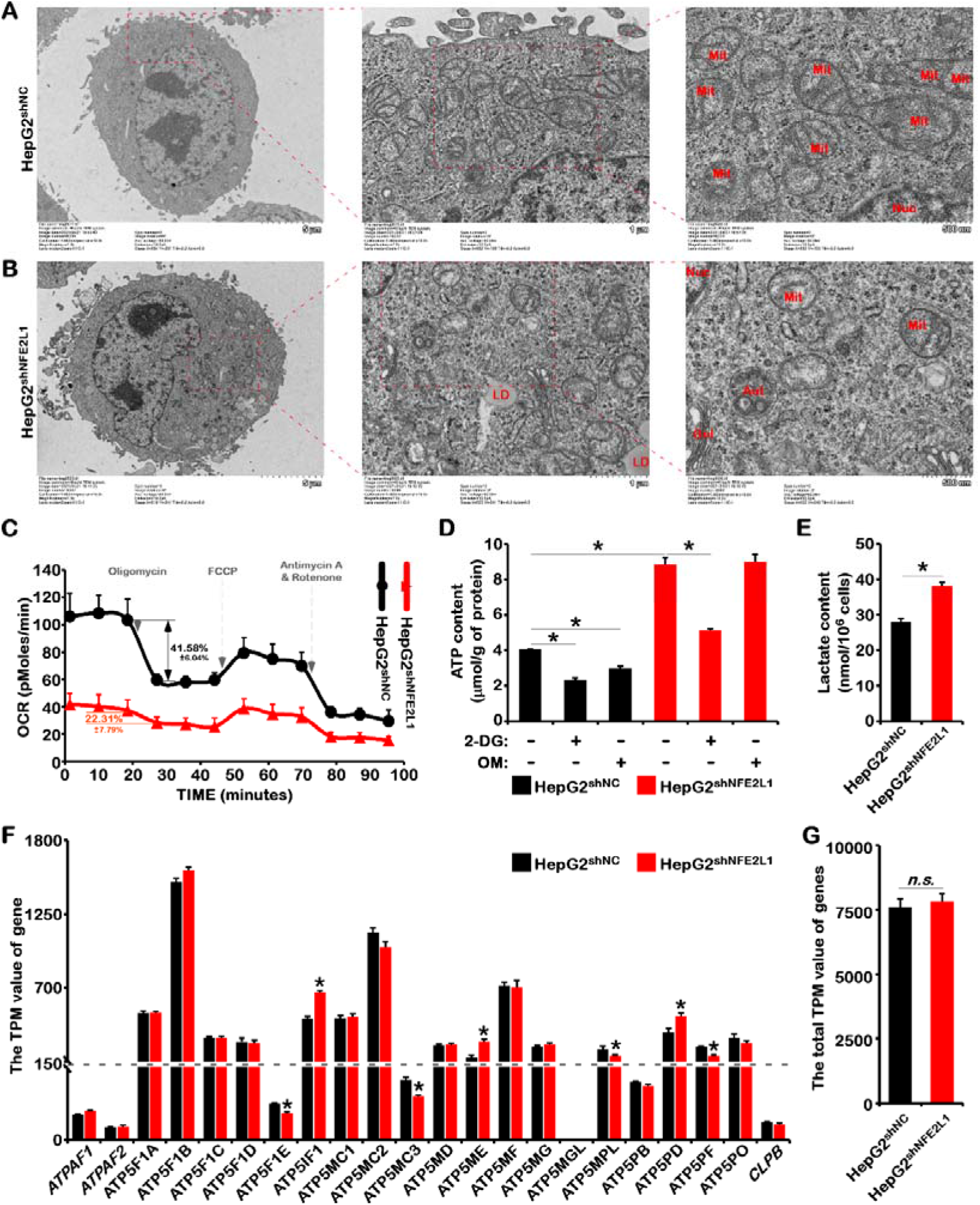
The effect of NFE2L1 knockdown in HepG2 cells on the morphology and function of mitochondria. The morphology of mitochondria in HepG2^shNC^ cells (**A**) and HepG2^shNFE2L1^ cells (**B**) were detected by electron microscope. (**C)** The oxygen consumption rate (OCR) of HepG2^shNC^ and HepG2^shNFE2L1^ cells were measured by Seahorse XF Cell Mito Stress Test Kit. (**D)** The HepG2^shNC^ and HepG2^shNFE2L1^ cells were treated with 2DG (20mM) or OM (10μM) for 12h, and the content of ATP in cells were detected by ATP assay kit. (**E)** The content of lactate in HepG2^shNC^ and HepG2^shNFE2L1^ cells were detected by Lactic Acid assay kit. (**F)** The TPM value of *ATPAF1*, *ATPAF2*, *ATP5F1A*, *ATP5F1B*, *ATP5F1C*, *ATP5F1D*, *ATP5F1E*, *ATP5IF1*, *ATP5MC1*, *ATP5MC2*, *ATP5MC3*, *ATP5MD*, *ATP5ME*, *ATP5MF*, *ATP5MG*, *ATP5MGL*, *ATP5MPL*, *ATP5PB*, *ATP5PD*, *ATP5PF*, *ATP5PO* and *CLPB* genes that related to ATP synthesis in mitochondria in HepG2^shNC^ and HepG2^shNFE2L1^ cells, date comes from transcriptome sequencing. (**G)** The total TPM value of genes in (**F**). n ≥ 3, ‘*’means *p* < 0.05, ‘*n.s.*’ means ‘no significant’.

By testing the oxygen consumption rate (OCR) of the cells, it was found that the oxygen consumption of the cells was reduced to about 40% of the control group after NFE2L1 knockdown (Figure 5C). The oxygen consumption of the control cells was reduced by 41.56%±6.04%, while in the NFE2L1 knockdown cells it was reduced by 22.31%±7.79% (Figure 5C) after adding OM, the oxidative phosphorylation inhibitor, indicating that the proportion of adenosine triphosphate (ATP) produced by mitochondrial respiration was significantly reduced after NFE2L1 knockdown. In the control group, the ATP content was reduced under treatment with 2DG or OM (Figure 5d). In addition, the lactate content significantly increased after NFE2L1 knockdown (Figure 5E). These results indicated that mitochondrial function was severely impaired after NFE2L1 knockdown.

In cells lacking NFE2L1 expression, although mitochondrial function was impaired and the oxidative phosphorylation process was inhibited, the ATP content increased more than twice (Figure 5D), meaning that compared with the control group, cells lacking NFE2L1 needed to consume dozens of times more glucose to produce these ATP through glycolysis. This might be the reason why lack of NFE2L1 led to increased cellular glucose uptake and cell death by glucose deprivation. Loss of NFE2L1 resulted in increased ATP in cells and the changed mitochondria morphology, suggesting the potential changes of ATP synthesis-related genes. Transcriptome data showed that the mRNA level of *ATP5IF1*, *ATP5ME* and *ATP5PD* genes significantly increased, and *ATP5F1E*, *ATP5MC3*, *ATP5MPL* and *ATP5PF* remarkably decreased after NFE2L1 knockdown (Figure 5F). No significant difference of genes related to ATP synthesis was observed in mitochondria (Figure 5G) despite the impacts of NFE2L1 deficiency in the characterization of mitochondria, indicating NFE2L1 might indirectly regulate the total amount of ATP. It is worth noting that NFE2L1 knockdown could still activate AMPK signal in the presence of high ATP concentration (Figure 1G), suggesting that the influence of NFE2L1 on AMPK signal may be through interfering with AMPK’s perception of ATP and adenosine monophosphate (AMP) levels in cells.

### 3.5. NFE2L1 serves as an energy-sensitive protein and directly inhibits the phosphorylation of AMPK by serine/threonine kinase 11 (STK11/LKB1)

Lack of NFE2L1 could disrupt AMPK signal which is the core energy metabolism regulation pathway [13–15]. Exploring the effect of NFE2L1 on AMPK signal is helpful to understand the influence of NFE2L1 on the metabolic network. To verify whether NFE2L1 is affected by ATP or AMP, exogenous ATP or AMP was used to treat HepG2 cells. The western blotting results suggested that the addition of exogenous ATP could increase the protein level of NFE2L1, while exogenous AMP showed the opposite effects (Figure 6A). This result was consistent with the response of NFE2L1 to glucose and serum (Figure 2), indicating that the response of NFE2L1 to the energy state might be due to ATP and AMP content in the cell. Subsequently, the effect of AMP on the phosphorylation of AMPK after NFE2L1 knockdown was tested, and the results indicated that NFE2L1 knockdown can reverse the activation of AMPK signal induced by AMP (Figure 6B,C). Interestingly, previous study showed NFE2L1 knockdown can reverse the activation of AMPK signal induced by MET [13]. These results were consistent, which showed the inhibition of NFE2L1 expression, indicating that activation of AMPK signaling by AMP and MET depended on the inhibitory effect of NFE2L1.

**Figure 6.**
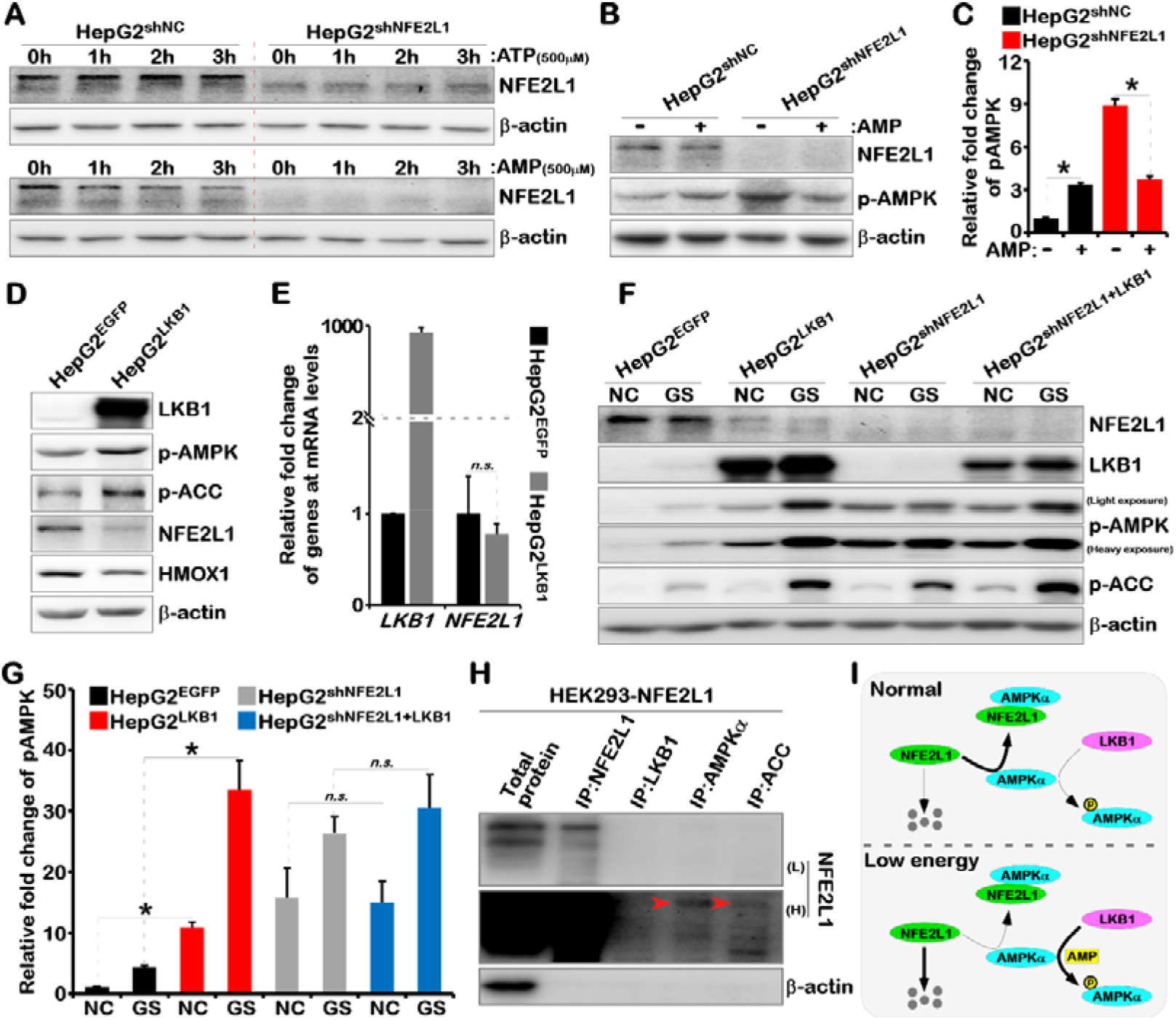
NFE2L1 disrupts the phosphorylation of AMPK by LKB1 by directly interacting with AMPK. **(A)** The HepG2^shNC^ and HepG2^shNFE2L1^ cells were treated with ATP (500μM) or AMP (500μM) for 1h, 2h, 3h, respectively. The expression of NFE2L1 and β-actin were detected by WB. (**B)** The HepG2^shNC^ and HepG2^shNFE2L1^ cells were treated with AMP (500μM) for 2h and the expression of NFE2L1, p-AMPK and β-actin were detected by WB. (**C)** Count and calculate the relative fold change of p-AMPK in (**B**). (**D)** The expression of LKB1, p-AMPK, p-ACC, NFE2L1, HO1 and β-actin in HepG2^EGFP^ and HepG2^LKB1^ cells were detected by WB. HepG2^EGFP^ and HepG2^LKB1^ cells were constructed by lentiviral infected. (**E)** The expression of *LKB1* and *NFE2L1* in HepG2^EGFP^ and HepG2^LKB1^ cells were detected by qPCR, with the β*-actin* was used as the internal control. (**F)** HepG2^EGFP^, HepG2^LKB1^, HepG2^shNC^ and HepG2^shNFE2L1+LKB1^ cells were treated with glucose starvation (GS) for 4h, and the expression of NFE2L1, LKB1, p-AMPK, p-ACC and β-actin were detected by WB. (**G)** Count and calculate the relative fold change of p-AMPK in (**F**). (**H)** The NFE2L1 overexpression plasmid *pLVX-NFE2L1-puro* transfected into HEK293T cells, and the total protein was collected with non-denatured lysate buffer after 48h. The specific antibodies of NFE2L1, LKB1, AMPK, and ACC protein were used for Co-IP experiments, then NFE2L1 and β-actin were detected by WB. ‘L’ means ‘Low exposure’ and ‘H’ means ‘Heavy exposure’. (**I)** The model diagram of NFE2L1 involved in regulating AMPK signal. n ≥ 3, ‘*’ means *p* < 0.05, ‘*n.s.*’ means ‘no significant’.

In the low-energy state, the ATP content in the cell decreases while the AMP content increases, which induces AMPK phosphorylation by LKB1 [35–37]. The activation of AMPK by LKB1 requires the co-localization of the two proteins in the plasma membrane. Here, our results showed that loss of NFE2L1, which is a membrane protein [38], inhibited AMPK signal activation induced by AMP, suggesting that NFE2L1 might be involved in the process of AMPK phosphorylation induced by LKB1.

In order to verify the effect of NFE2L1 on LKB1-induced AMPK phosphorylation, LKB1 overexpressing and NFE2L1 knockdown HepG2 cells were constructed. Unexpectedly, in HepG2 cells, overexpression of LKB1 could activate AMPK signal and significantly inhibit the protein level of NFE2L1, but showed no obvious effects on the mRNA level of *NFE2L1* (Figure 6D,E), indicating the coincidence of NFE2L1 reduction and AMPK phosphorylation increase. In cells with NFE2L1 deficiency, overexpression of LKB1 could not further increase AMPK phosphorylation (Figure 6F,G). Glucose starvation (GS) experiment results showed that glucose starvation for 2h significantly increased AMPK phosphorylation, and GS could still further increase AMPK phosphorylation after LKB1 overexpression (Figure 6F,G). The cumulative effects of LKB1 overexpression and GS treatment on AMPK phosphorylation levels suggested that the response of AMPK to low-energy states might not completely depend on LKB1. In the NFE2L1 knockdown cells, with the increase of the basal AMPK phosphorylation, although GS treatment further increased AMPK phosphorylation, the cumulative effect of LKB1 overexpression and GS treatment disappeared (Figure 6F,G), suggesting that the effects of NFE2L1 on AMPK signal might mainly relied on LKB1.

Subsequently, protein immunoprecipitation experiments were utilized to explore the action mode of NFE2L1 on LKB1-AMPK. The results showed that AMPK could interact with NFE2L1 protein rather than LKB1 (Figure 6H), suggesting that NFE2L1 might directly bind to AMPK and inhibit LKB1’s phosphorylation. In addition, ACC could also interact with NFE2L1 (Figure 6H). Combined with the negative correlation between phosphorylated ACC and NFE2L1 (Figure 2A,B), we could speculate that NFE2L1 may regulate lipid metabolism by directly binding to ACC and inhibiting its phosphorylation.

## 4. Discussion

The rational use of nutrients is a necessary prerequisite for the survival of multicellular organisms. In this process, the perception and feedback of the energy state of cells is crucial. NFE2L1 has been considered to play an important role in regulating system redox homeostasis, carbohydrate and lipid metabolism homeostasis, embryonic development and cancer [4]. The transcription factor NFE2L1 systematically regulates the transcription of antioxidant-related genes through the recognition of antioxidant/electrophile-response element (ARE/EPRE) [39]. However, the molecular mechanism of metabolism-related phenotypes caused by NFE2L1 overexpression or knockout is still elusive. Identification of the molecular mechanism of NFE2L1 in regulating substance and energy metabolism will deepen the recognition of the molecular and physiological functions of NFE2L1, which could benefit for the treatment of obesity, diabetes, non-alcoholic fatty liver disease and other systemic metabolic diseases.

NFE2L1 is essential for maintaining intracellular redox homeostasis which is a prerequisite for cells to maintain normal metabolic processes. Loss of NFE2L1 can not only directly increase ROS levels, but also induce strong oxidative stress to cause the death of embryos [1]. In the NFE2L1 knockdown cell line, glucose deprivation could induce cell death, which might be related to the imbalance of redox homeostasis (Figure 1). These results were consistent with previous studies [12]. NFE2L1 could negatively regulate glucose uptake (Figure 3). These results demonstrated the dual functions of NFE2L1 as the sensor and the regulator of glucose homeostasis.

The increased glucose uptake caused by NFE2L1 knockdown might be related to the increased of GLUT3 (Figure 3). Previous reports mostly focused on studying the relationship between NFE2L1 and GLUT1, GLUT2, GLUT4 respectively [6,7,12]. However, the interaction of NFE2L1 with GLUT3, the transporter with the highest affinity for glucose [40], has not been explored. Recently, more and more studies have shown that GLUT3 plays a vital role in regulating glycolysis, cancer stem cell survival, and cancer cell metastasis [41–43]. In addition, GLUT3, which can be activated by AMPK signals [44], is mainly expressed in tissues with high glucose requirements and poor glucose microenvironment, such as the brain [45]. Here, we found that knockdown of NFE2L1 could significantly increase GLUT3 expression, which might be attributed to AMPK activation. NFE2L1 specific knockout led to neurodegenerative diseases in the nervous system [46] and metabolic disorders and hepatocellular carcinogenesis in liver tissue [2]. However, the expression of GLUT3 gene was decreased in NFE2L1 knockdown cells, making it not easy to identify the underlying molecular mechanism. It is worth mentioning that the glycosylation and deglycosylation of NFE2L1 are necessary to activate its transcription factor activity [19]. Meanwhile, the glycation of NFE2L1 strictly depended on the glycolysis process (Figure 2E–H). These results suggested that the NFE2L1 functioned as a key factor in mediating the glucose homeostasis and redox homeostasis in cells.

Studies have shown that NFE2L1 is sensitive to nutrients in the environment [23]. However, the growth of cells lacking NFE2L1 depended almost solely on glucose. By detecting the key rate-limiting enzymes involved in glucose metabolism and the proteomic of glucose metabolites, we found that loss of NFE2L1 significantly inhibited the oxidative phosphorylation process and enhanced the glycolysis and gluconeogenesis process (Figure 4). Obviously, NFE2L1 deficiency in HepG2 cells triggered and enhanced the Warburg effect, which might be the cause of spontaneous NASH that could be finally developed into HCC in the liver of *NFE2L1* knockout mice [2].

Overexpression of NFE2L1 can reverse the mitochondrial damage caused by N-glycanase 1 (NGLY1) knockout [25], suggesting that NFE2L1 is essential for the maintenance of mitochondrial morphology and function. Here, we also observed that knockdown of NFE2L1 can cause mitochondrial damage (Figure 5). However, the relationship between the Warburg effect and mitochondrial function damage caused by NFE2L1 deficiency is still hard to determine. Once mitochondria, as the most important organelle for the production of ATP, are damaged, cells may need to consume more glucose to produce ATP through glycolysis. As the results showed, glucose uptake capacity increased after NFE2L1 knockdown. And after deprivation of glucose, cells with NFE2L1 knockdown died. Strikingly, The ATP content in HepG2^shNFE2L1^ cells was significantly higher than that in HepG2^shNC^ cells (Figure 5D), which is consistent with the results of Zhu et al. [12]. OM, the inhibitor of oxidative phosphorylation, could hardly inhibit ATP production in HepG2^shNFE2L1^ cells, indicating that these ATP were rarely produced from the oxidative phosphorylation process (Figure 5D). In short, HepG2^shNFE2L1^ cells need to maintain a higher level of ATP content with lower ATP production efficiency. This may be the reason of the dependence of the growth of HepG2^shNFE2L1^ cells on glucose.

AMPK functions as the core regulator of energy metabolism to sense the levels of ATP and AMP. The binding of AMP with AMPK is necessary for the activation of AMPK by phosphorylation [37]. In the low energy state, the increase of AMP caused by the consumption of ATP can activate AMPK signals to promote catabolism and inhibit synthesis reactions which could finally produce more ATP [15]. The increase of phosphorylated AMPK level after NFE2L1 knockdown (Figure 1E) might be responsible for the reprogramming of glucose metabolism and the increase of ATP (Figures 3, 4). Under normal circumstances, high levels of ATP could inhibit AMPK signaling to maintain energy homeostasis. Obviously, NFE2L1 knockdown destroyed the energy homeostasis maintained by this mechanism. High ATP content and highly activated AMPK signals coexist in the HepG2^shNFE2L1^ cells, suggesting the problem of the negative feedback regulation of AMPK signals.

In the HepG2^shNFE2L1^ cells, AMP lost the ability of increasing the level of AMPK phosphorylation (Figure 6B,C), which once again confirmed that NFE2L1 might be involved in AMPK signal perception of ATP and AMP. In other words, AMP binding is not necessary for the phosphorylation of AMPK in the absence of NFE2L1. Interestingly, various methods that can improve AMPK phosphorylation can also lead to decreased NFE2L1, including FBS starvation [23], glucose starvation [13], MET treatment [13], AMP treatment (Figure 6A), and LKB1 overexpression (Figure 6D). On the contrary, treatments with EGF and insulin [23] that can inhibit AMPK phosphorylation are accompanied by the increase of NFE2L1. In addition, NFE2L1 could interact with AMPK protein (Figure 6H). All these results revealed that NFE2L1 could bind to AMPK in the cytoplasm to inhibit the phosphorylation of AMPK in addition to the function as a transcription factor in the nucleus. In a low-energy state, the binding of NFE2L1 with AMPK reduced due to the lower expression level of NFE2L1 protein, which decreased the inhibition of NFE2L1 to AMPK activity (Figure 6I). Also, AMPK could bind with NFE2L1 and inhibit the degradation of NFE2L1. However, the sequential relationship between the decrease of NFE2L1 protein and the increase of AMPK phosphorylation is still unknown. Whether the effects of AMP/ATP and LKB1 on AMPK depend on NFE2L1 requires to be explored.

As an environment-sensitive protein, NFE2L1 could be changed in response to various stresses even the environmental temperature changes [4,47]. NFE2L1 could function as a transcription factor to directly regulate the transcription and expression of thousands of genes involved in almost all biological processes [32,48]. In addition, with the half-life of about 0.5h, NFE2L1 protein is always in the process of dynamic synthesis and degradation [13]. These characteristics make NFE2L1 own the ability of mediating the interaction and dialogue between the internal homeostasis and the external environment. In this study, we discovered for the first time that NFE2L1 could directly regulate the activity of AMPK signaling in a way independent of the function of transcription factor. The findings could deepen the understanding of the molecular mechanism of NFE2L1 in regulating material and energy metabolism, and redox homeostasis, which is valuable for targeting NFE2L1 to prevent or treat systemic diseases.

## 5. Conclusions

In conclusion, the results of glucose starvation, glucose uptake, selective inhibition of glucose metabolism and protein deglycosylation experiments suggested that NFE2L1 could act as the sensor and the regulator of glucose homeostasis. Transcriptome, metabolome, seahorse and electron microscopy results revealed that impaired NFE2L1 caused reprogram of glucose metabolism, damaged the mitochondrial and aggravated the Warburg effect. And the energy signal transmission and Co-IP experiments found that NFE2L1 could sense the energy state and regulate AMPK signaling pathway by directly interact with AMPK protein. The novel AMP/NFE2L1/AMPK signaling pathway that discovered in this study may be the core mechanism of NFE2L1 in metabolism and metabolic diseases.

## Supplementary Materials

The following are available online at xxx. **Table S1.** The key resources in this study. **Table S2.** The transcriptome sequencing data of HepG2^shNC^ and HepG2^shNFE2L1^ cell lines.

## Author Contributions

LQ and YQ conceived the research and designed the experiments; QY, WZ, YX, PL, XZ, HN, RS, SG, and YC conducted the experiments, acquired and analyzed the data with the critical assistance from WJZ, YW, GL, ZC, YR, YX, YG and YZ; all authors contribute to writing the manuscript; WZ, XZ, YX, YZ and LQ revised the manuscript. All authors read and approved the final manuscript.

## Funding

This work was supported by the Postdoctoral Research Grant in Henan Province (201901006, 201902006), the China Postdoctoral Science Foundation (2020M672286), the Program for Innovative Talents of Science and Technology in Henan Province (18HASTIT042), Science Foundation for Excellent Young Scholars in Henan (202300410358) and National Natural Science Foundation of China (U1904147, U20A20369, 81822043).

## Data Availability Statement

The data presented in this study are available in this manuscript.

## Conflicts of Interest

The authors declare no conflict of interests.

## Notes

### Competing Interest Statement

The authors have declared no competing interest.

