## Supplementary Tables for "Dysfunction of an energy sensor NFE2L1 triggers uncontrollable AMPK signal and glucose metabolism reprogramming": Supplementary Table 1. The key resources.docx

**Table S1.** The key resources

| REAGENT or RESOURCE | SOURCE | | IDENTIFIER |
| --- | --- | --- | --- |
| Cell Lines | | | |
| HepG2 | Cell bank of the Chinese Academy of Sciences | | TCHu72 |
| HepG2^shNC^ | Laboratory preservation | | [1] |
| HepG2^shNFE2L1^ | Laboratory preservation | | [1] |
| HepG2^EGFP^ | this manuscript | | N/A |
| HepG2^LKB1^ | this manuscript | | N/A |
| HepG2^shNFE2L1+EGFP^ | this manuscript | | N/A |
| HepG2^shNFE2L1+LKB1^ | this manuscript | | N/A |
| HEK293T | Laboratory preservation | | N/A |
| Chemicals | | | |
| WZB117 | APExBIO | | A8244 |
| Glucose (GLU) | Sangon Biotech | | A600219 |
| D-(-)-Fructose (FRU) | Sangon Biotech | | A600213 |
| D-(+)-Mannose (MAN) | Sangon Biotech | | A600554 |
| Fructose 1,6-bisphosphate (FBP) | Sangon Biotech | | A600468 |
| pyruvate (PYR) | Sangon Biotech | | A100342 |
| D-(+)-Galactose (GAL) | Sangon Biotech | | A600215 |
| D-(-)-Ribose (RIB) | Sangon Biotech | | A610475 |
| Endo H | New England Biolabs | | P0702S |
| Oligomycin (OM) | Sangon Biotech | | A606700 |
| 2-Deoxy-D-glucose (2-DG) | APExBIO | | B1027 |
| 2-NBDG | APExBIO | | B6035 |
| ATP | Meilunbio | | 34369-07-8 |
| AMP | Sigma Aldrich | | A1752-1G |
| Antibodies | | | |
| p-ACC (Phospho-Ser79) | Sangon Biotech | | D155180 |
| ACC | Sangon Biotech | | D155300 |
| p-AMPK (Phospho-Thr183/Thr172) | Sangon Biotech | | D151212 |
| AMPK | Sangon Biotech | | D261273 |
| p-mTOR | CST | | #5536 |
| p-S6K1 | Sangon Biotech | | D151473 |
| KEAP1 | Sangon Biotech | | D260984 |
| PTGS2 | Abcam | | ab62331 |
| HK1 | Sangon Biotech | | D221854 |
| PFKL | Sangon Biotech | | D222865 |
| PKM | Sangon Biotech | | D162511 |
| PC | Sangon Biotech | | D221117 |
| IDH1 | Sangon Biotech | | D221821 |
| IDH2 | Sangon Biotech | | D222530 |
| PCK2 | Sangon Biotech | | D222686 |
| GLUT1 | Abcam | | ab115730 |
| GLUT2 | Abcam | | ab192599 |
| GLUT3 | Abcam | | ab191071 |
| GLUT4 | Bioss | | bs-0384R |
| LKB1 | Abcam | | ab199970 |
| HMOX1 | Sangon Biotech | | D220756 |
| NFE2L1 | CST | | #8052 |
| NFE2L2 | Abcam | | ab62352 |
| β-actin | ZSGB-BIO | | TA-09 |
| Oligonucleotides for qPCR | | | |
| GLUT1 FW | Sangon Biotech | | GCAGCTACCCTGGATGTCCTATCT |
| GLUT1 REV | Sangon Biotech | | CTGGAAGCACATGCCCACAATGA |
| GLUT2 FW | Sangon Biotech | | AGCTACCGACAGCCTATTCTAGTG |
| GLUT2 REV | Sangon Biotech | | CAAACATCCCACTCATTCCAAT |
| GLUT3 FW | Sangon Biotech | | CCAACTTCCTAGTCGGATTGCT |
| GLUT3 REV | Sangon Biotech | | GTGGTCTCCTTAGCAGGCTCGAT |
| GLUT4 FW | Sangon Biotech | | TTCCAGTATGTTGCGGAGGCTAT |
| GLUT4 REV | Sangon Biotech | | CTGGGTTTCACCTCCTGCTCTA |
| GLUT5 FW | Sangon Biotech | | GTGGTCTGTAACCGTGTCCATGTT |
| GLUT5 REV | Sangon Biotech | | AGCTCAAATGATGTGGCGACTCT |
| GLUT6 FW | Sangon Biotech | | CGGTGTACGTGTCTGAGATTGCT |
| GLUT6 REV | Sangon Biotech | | ATGAAGCTGAGCAGCAGGATCAT |
| GLUT7 FW | Sangon Biotech | | CCGCACAAGGTCTTCAAGTCAT |
| GLUT7 REV | Sangon Biotech | | GGCAAAGATGTTGTTGATCAGCA |
| GLUT8 FW | Sangon Biotech | | GTCTACATCTCCGAAATCGCCTA |
| GLUT8 REV | Sangon Biotech | | GAAGCACATGAGAAGCAGCATGA |
| GLUT9 FW | Sangon Biotech | | TCCCATACGTCACCTTGAGTACA |
| GLUT9 REV | Sangon Biotech | | GAAGAACTCACCAGTCAAGATGAAC |
| NFE2L1 FW | Sangon Biotech | | CACTCCCATCAATCAGAATGTCA |
| NFE2L1 REV | Sangon Biotech | | AGGTTGGTGGAGCCGAAGGT |
| HMOX1 FW | Sangon Biotech | | GCCAGCAACAAAGTGCAAGAT |
| HMOX1 REV | Sangon Biotech | | GGTAAGGAAGCCAGCCAAGAGA |
| PTGS1 FW | Sangon Biotech | | CGCCAGTGAATCCCTGTTGTT |
| PTGS1 REV | Sangon Biotech | | AAGGTGGCATTGACAAACTCC |
| PTGS2 FW | Sangon Biotech | | AAGTCCCTGAGCATCTACGGTTT |
| PTGS2 REV | Sangon Biotech | | GTTGTGTTCCCTCAGCCAGATT |
| HK1 FW | Sangon Biotech | | CACATGGAGTCCGAGGTTTATG |
| HK1 REV | Sangon Biotech | | CGTGAATCCCACAGGTAACTTC |
| HK2 FW | Sangon Biotech | | GAGCCACCACTCACCCTACT |
| HK2 REV | Sangon Biotech | | CCAGGCATTCGGCAATGTG |
| PFKL FW | Sangon Biotech | | GGCTTCGACACCCGTGTAA |
| PFKL REV | Sangon Biotech | | CGTCAAACCTCTTGTCATCCA |
| PFKM FW | Sangon Biotech | | GGTGCCCGTGTCTTCTTTGTC |
| PFKM REV | Sangon Biotech | | CAGCTCGGAGTCGTCCTTCTC |
| PKM FW | Sangon Biotech | | ATGTCGAAGCCCCATAGTGAA |
| PKM REV | Sangon Biotech | | TGGGTGGTGAATCAATGTCCA |
| PC FW | Sangon Biotech | | CTGAATACTCGCCTCTTCCTGC |
| PC REV | Sangon Biotech | | CTTCATGGCCTGGGTGTCCTTG |
| DLD FW | Sangon Biotech | | GAAATGTCCGAAGTTCGCTTGA |
| DLD REV | Sangon Biotech | | TCAGCTTTCGTAGCAGTGACT |
| CS FW | Sangon Biotech | | TGCTTCCTCCACGAATTTGAAA |
| CS REV | Sangon Biotech | | CCACCATACATCATGTCCACAG |
| IDH1 FW | Sangon Biotech | | TGTGGTAGAGATGCAAGGAGA |
| IDH1 REV | Sangon Biotech | | TTGGTGACTTGGTCGTTGGTG |
| IDH2 FW | Sangon Biotech | | TGGCAGTGGTGTCAAGGAGTG |
| IDH2 REV | Sangon Biotech | | GCCCATCGTAGGCTTTCAGTA |
| IDH3A FW | Sangon Biotech | | TCACCCATCTATGAATTTACTGCTG |
| IDH3A REV | Sangon Biotech | | GATACTCTGCACGACTCCATCAAC |
| IDH3B FW | Sangon Biotech | | GAGCCAAGTCTCAGCGGAT |
| IDH3B REV | Sangon Biotech | | GGGCATCACAAGCACATCAA |
| IDH3G FW | Sangon Biotech | | GACCCGGCACAAGGACATAG |
| IDH3G REV | Sangon Biotech | | GCTTGAAGGCATACTCGGCAA |
| OGDH FW | Sangon Biotech | | TAGAAGGCTGCGAGGTACTGA |
| OGDH REV | Sangon Biotech | | CGATTGATCCTGCGGTGATA |
| G6PC3 FW | Sangon Biotech | | CGAGGCGCTACAGAACCAG |
| G6PC3 REV | Sangon Biotech | | CACTCGGTGATGAGGCTGAT |
| FBP1 FW | Sangon Biotech | | CTCTATGGCATTGCTGGTTCTAC |
| FBP1 REV | Sangon Biotech | | GGTTCCACTATGATGGCGTGT |
| PCK2 FW | Sangon Biotech | | AGTAGAGAGCAAGACGGTGAT |
| PCK2 REV | Sangon Biotech | | TGCTGAATGGAAGCACATACAT |
| LKB1 FW | Sangon Biotech | | CGGCTTCAAGGTGGACATCTGGT |
| LKB1 REV | Sangon Biotech | | GTTCGTACTCAAGCATCCCTTTCAG |
| β-actin FW | Sangon Biotech | | CATGTACGTTGCTATCCAGGC |
| β-actin REV | Sangon Biotech | | CTCCTTAATGTCACGCACGAT |
| Oligonucleotides for construct plasmid | | | |
| LKB1 FW | Sangon Biotech | CGCCTCGAGATGGAGGTGGTGGA | |
| LKB1 REV | Sangon Biotech | CGCGAATTCTGCTGCTTGCAGGC | |
| NFE2L1 FW | Sangon Biotech | CTCGAGGAATTCATGCTTTCTCTGAAGAAATAC | |
| NFE2L1 REV | Sangon Biotech | GGCCGCTCTAGATCACTTTCTCCGGTCCTTTG | |
| Plasmid vector | | | |
| *pLVX-puro* | Biofeng | | PT4002-5 |
| *pMD2.G* | addgene | | 12259 |
| *psPAX2* | addgene | | 12260 |
| *pLVX-EGFP-puro* | this manuscript | | N/A |
| *pLVX-LKB1::EGFP-puro* | this manuscript | | N/A |
| *pLVX-NFE2L1-puro* | this manuscript | | [1] |
| Software | | | |
| Canvas X | Canvas GFX, Inc. | | https://www.canvasgfx.com/ |
| Chromas 2.4.1 | [Technelysium Pty Ltd](https://www.baidu.com/link?url=Z4b5CJjCWWFkUOLD-A5XGqStVl1TGC7fGWGoe8Z7MSAzDX5MFgCT27KZTsARjy9N&wd=&eqid=e2341667000303b1000000055b6be0a6) | | http://technelysium.com.au/wp/chromas/ |
| Excel | Microsoft | | https://www.microsoft.com/ |
| Quantity One 4.5.2 | Bio-Rad | | https://www.bio-rad.com/ |
| Seahorse Analytics | Agilent | | https://www.agilent.com.cn/ |
| Primer Premier 5 | PREMIER Biosoft International | | https://www.PremierBiosoft.com/ |
| Kits | | | |
| Lipofectamine® 3000 Transfection Kit | Invitrogen | | L3000-015 |
| TIANpuro Midi Plasmid Kit | Tiangen Biotech | | DP107 |
| Reactive Oxygen Species Assay Kit | Beyotime | | S0033 |
| ATP Assay Kit | Beyotime | | S0026 |
| Lactic Acid assay kit | Nanjing Jiancheng Bioengineering Institute | | A019-2-1 |
| Seahorse XF Cell Mito Stress Test Kit | Agilent | | 103015-100 |
| Cell Counting Kit-8 | Beyotime | | C0039 |
| RNAsimple Total RNA Kit | Tiangen Biotech | | DP419 |
| Revert Aid First Strand Synthesis Kit | Thermo | | K1622 |
| GoTaq® qPCR Master Mix | Promega | | A6001 |

1.	Gou, S.; Qiu, L.; Yang, Q.; Li, P.; Zhou, X.; Sun, Y.; Zhou, X.; Zhao, W.; Zhai, W.; Li, G.*, et al.* Metformin leads to accumulation of reactive oxygen species by inhibiting the nfe2l1 expression in human hepatocellular carcinoma cells. *Toxicol Appl Pharmacol* **2021**, *420*, 115523.
